# Predicting Protein-RNA Binding Affinity Changes via Spatial Coupling-Aware State Space Modeling

**DOI:** 10.64898/2026.08.23.745486

**Authors:** Rui Chen, Xiaoyu Huang, Huasen Jiang, Wenjian Ma, Xiangpeng Bi, Zhiqiang Wei, Jie Nie, Shugang Zhang

## Abstract

Accurately predicting the effects of mutations on protein-RNA binding is crucial for elucidating disease mechanisms. Yet, exhaustively exploring the space of all possible variants is prohibitively expensive, motivating computational methods that can quantify mutation-induced changes in binding affinity (aka ΔΔG) accurately and efficiently. We present **iSCALE**, an interpretable and generalizable deep learning method that adopts an **i**mplicit **S**patial **C**oupling-**A**ware **L**igand **E**ncoding strategy to predict mutation-induced binding affinity changes. By injecting this implicit multiscale encoding scheme into a bidirectional state space modeling architecture, iSCALE learns a generalizable multiscale coupling pattern that achieves superior performances on not only the protein-RNA binding ΔΔG, but also the protein stability ΔΔG and protein-protein binding ΔΔG predictions. Detailed analyses demonstrate that the model attention scores align well with structural characteristics. In addition, iSCALE shows good discriminative ability when predicting close samples such as complexes of same mutation but with different ligands or the same complex but with different mutation sites. In summary, iSCALE serves as an effective in silico tool for large-scale protein-RNA binding ΔΔG prediction, which pushes the border of understanding in mutation-induced pathological outcomes.

## I. Introduction

Protein-RNA interactions are central to post-transcriptional gene regulation, mediating crucial biological processes like RNA splicing, mRNA stability, translational control [1], [2]. The human genome encodes more than 1,500 human RNA-binding proteins containing approximately 600 structurally distinct RNA binding domains [1], [3] [4], underscoring the vast landscape and evolutionary significance of protein-RNA interactions. Disruption of these interactions has been linked to diverse human diseases including cancer and neurodegeneration, and other genetic disorder [5]. Quantifying mutation-induced changes in protein-RNA binding affinity (i.e., ΔΔG) therefore provides a mechanistic basis for interpreting pathogenic variants and prioritizing functional mutations.

Despite their biological importance, the functional consequences of mutations remain difficult to comprehensively deciphered due to the numerous variants and the weak mutation signals. For instance, a typical protein of 500 residues would give rise to nearly 10,000 potential single-site substitutions, before considering the exponentially expanding combinations of multi-site mutations, as well as the highly ligand-specific outcomes for different RNAs upon binding. Exhaustive experimental characterization of these variants is therefore impractical, which creates a pressing need for computational methods that can learn sequence-structure-function relationships from available data and predict mutation-induced binding ΔΔG.

Early computational approaches for modelling mutation-induced changes in protein-RNA binding primarily relied on energy-based tools, exemplified by FoldX [6] and machine learning extensions like mmCSM-NA [7], PremPRI [8], and PRA-MutPred [9]. These methods rely on predefined empirical energy functions and are computationally intensive upon large-scale mutational screening. Therefore, there is increasing interest in deep learning methods as a data-driven paradigm. Existing approaches are categorized into sequence-based [10] and structure-based models [11], [12].

Sequence-based methods such as PRITrans [13] and DeePNAP [14] integrate protein language-model embeddings [15], [16], nucleotide or k-mer encodings and attention-based fusion to predict protein-RNA interactions from primary sequences alone. However, deep learning architectures tailored to sequence data are not ideally matched to the mutation-effect prediction. For example, Transformer and its derived language models are powerful in capturing long-range sequence context, but meanwhile fall short in capturing subtle mutation-induced signals[17]. Furthermore, functionally important mutations are often accompanied by conformational changes[18], sequence-only models apparently lack an explicit description of the structural aspects.

Structure-based deep learning methods address this limitation by representing complexes as residue-level or atom-level graphs [10], which are further enriched with geometric features [19]. Representative geometric learning architectures such as Geometric Vector Perceptrons (GVP) [20] and Equivariant Graph Neural Networks (EGNN) [21] capture structural details such as directional information and distances in a principled manner, allowing models to focus on mutation-centered local geometric context. A recent mutation-focused model MuToN describes protein structures and binding interface via structural encoders, and leverages geometric message passing to quantify how specific residue substitutions perturb local interaction patterns [22]. Structure-based models have improved the representation of protein-RNA recognition, but they also introduce a different set of challenges. Explicit modelling of fine-grained and atom-level geometric features can make models sensitive to structural noise, encouraging overfitting rather than learning a generalizable interaction pattern. In addition, these models are computationally heavier, architecturally more complex, and often rely on expert-defined structural descriptors.

To address these limitations, we introduce an interpretable and generalizable deep learning-based method named iSCALE, which adopts an **i**mplicit **S**patial **C**oupling-**A**ware **L**igand **E**ncoding strategy to predict mutation-induced binding affinity changes. Its core idea is to model the interaction geometry implicitly rather than using an explicit way like previous geometric learning methods. More specifically, iSCALE adopts a novel multiscale feature encoder to embed RNA information implicitly into protein residue features. This particular encoder follows a coarse-grained way and does not overly depict geometric details, thereby avoiding overfitting to detailed geometric patterns. After that, iSCALE employs a state space modeling module along the wild-type and mutant protein sequences to model long-range dependencies and propagate mutation effects. The above two modules form a sequence-structure coupled model architecture that being trained to predict ΔΔG. Simultaneously, an auxiliary reconstruction task is also introduced to enhance structural awareness.

Comprehensive evaluations and analyses reveal that iSCALE consistently surpassed existing methods on the benchmark protein-RNA binding ΔΔG prediction dataset, regardless of standard settings, reduced training data, or more stringent PDB-limited scenarios. The multiscale feature encoder, as a critical finding in this work, presents impressive universal applicability across baselines and tasks. It also aligns well with experimentally observed coupling patterns. Further exploration validates the ability of iSCALE to discriminate ligand-dependent effects of same mutations and distinct mutation-specific destabilizing effects within the same complex. In summary, iSCALE offers a unified and efficient methodology for predicting ΔΔG upon mutations. Its characterized multiscale distribution features reflect inherent biological principles shared by diverse molecular interactions.

## II. Results

### 2.1 Overview of iSCALE

iSCALE is a deep learning-based method for predicting mutation-induced protein-RNA binding affinity changes. The overall architecture of iSCALE (Fig. 1) consists of three modules: (1) a multiscale feature encoder module for describing input protein and ligand pairs. (2) a state space modeling module for learning sequence features. (3) a dual-task prediction module for jointly optimizing the main ΔΔG prediction task and an auxiliary spatial coupling distribution prediction task.

**Fig. 1.**
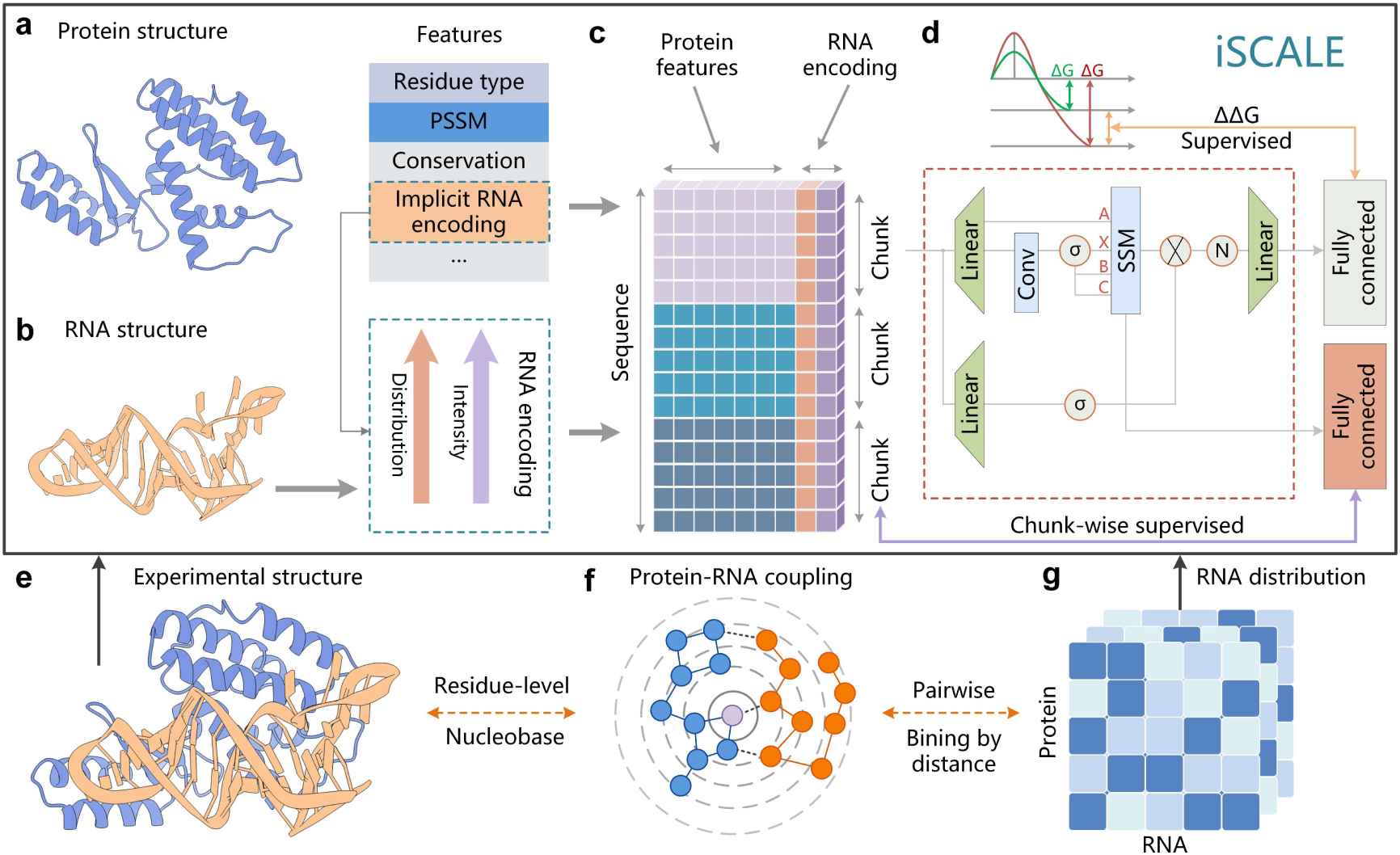
| Architecture of iSCALE. The framework predicts mutation-induced changes in protein–RNA binding affinity (ΔΔG) from protein sequence features augmented with multiscale structural information derived from the protein–RNA complex. **(a)** Protein residue features. Each residue is represented by amino acid residue type, position-specific scoring matrix (PSSM), conservation score, and an implicit structure component encoding the spatial relationship with the RNA ligand. **(b)** RNA structural information is implicitly encoded as two features appended to each protein residue, i.e., spatial coupling intensity and spatial coupling distribution. **(c)** The RNA-structure-encoded protein feature matrix is divided into non-overlapping chunks of 32 residues. Within each chunk, iSCALE performs structured masked attention inherited from SSD [23] to model the fine-grained pairwise interactions among residues. The chunk-level hidden state is in turn propagated across chunks via a linear recurrence to model long-range dependencies. **(d)** Dual-task prediction module. The SSD layer indicated by the red dashed box takes chunk-wise features as inputs. The learned representations are projected via fully connected layers into ΔΔG for prediction. Meanwhile, the hidden chunk expressions are fetched to predict the spatial coupling distribution, forming an auxiliary chunk-wise supervision task. **(e)** Three-dimensional structure of a protein– RNA complex. **(f)** For each protein residue (blue circles), pairwise distances to all RNA nucleotides (orange circles) are computed and assigned to bins defined by distance thresholds (concentric rings). **(g)** The resulting protein-centric distribution matrix: each row corresponds to one protein residue, each column to one distance bin, with cell intensity encoding the proportion of RNA nucleotides falling within the corresponding distance range.

The multiscale feature encoder module takes the protein-RNA complex as inputs (Fig. 1a-b). For the protein, the model adopts a residue-level descriptor consisting of amino acid type, PSSM, and conservation features. Besides, the residue-level descriptor also incorporates an implicit structure component describing the multiscale structural information of RNA ligand. The component takes forms of either the coupling intensity defined as a value that inversely proportional to the distance from a residue to its nearest nucleotide, or a distribution vector denoting the multiscale distribution of RNA nucleotides in terms of their distance from the residue (“Methods”). The RNA information is only encoded here into residue features without any additional modeling. Subsequently, the state space modeling module (Fig. 1c-d) takes the RNA-embedded protein sequence descriptor as inputs and models the long-range dependencies. Each sequence is divided into functional chunks of 32-residue in length. iSCALE performs linear attention within each chunk and propagates states across chunks. The wild-type and mutant sequences are concatenated and we adopt bidirectional processing to model the forward and reverse mutations. Finally, the dual-task prediction module (Fig. 1d-g) takes the learned sequence representation and predicts mutation-induced ΔΔG. The multiscale distribution of RNA nucleotides is utilized as an auxiliary task. Residue-level distributions are re-organized into chunk-wise labels to avoid inefficient and overly granular learning. As a biological-inspired design, the auxiliary task forces the model to reconstruct the multiscale conformational coupling information from the sequence space.

### 2.2 iSCALE enables accurate protein-RNA binding ΔΔG prediction by modeling multiscale conformational coupling

To evaluate iSCALE’s performance in predicting protein-RNA binding ΔΔG upon mutations, we curated a dataset containing 394 experimentally validated missense mutations covering 78 different protein-RNA complexes. We compared our model’s performance against eight existing models (Fig. 2a), including a variant of iSCALE that without auxiliary coupling prediction task (aka iSCALE*). To demonstrate that the effectiveness of implicit RNA structural information generalizes to different models, we append the two features, i.e., *spatial coupling intensity* and *spatial coupling distribution* to all models. It can be observed the distribution feature benefits almost all models (see the Distribution column). Specifically, PCC of graph-based models like GCN, DGCNN, etc., improved from 0.793–0.80 to 0.818–0.824, and Transformer-based methods improved from 0.928–0.941 to 0.942–0.948. In contrast, the coupling intensity feature provided smaller improvements or negative gains (see the Intensity column), and combining these two features does not show significant advantages compared to using the distribution feature alone. The performance difference between two features is expected, because the coupling intensity reflects only the nearest nucleotide while the distribution feature encodes information of all nucleotides via a multiscale way. Accordingly, the spatial coupling distribution feature is finalized in iSCALE, and it achieved the highest PCC of 0.969 among all existing models. The improvement of iSCALE over iSCALE* indicates that the auxiliary coupling prediction task provides additional structural supervision beyond what is available through input features.

**Fig. 2.**
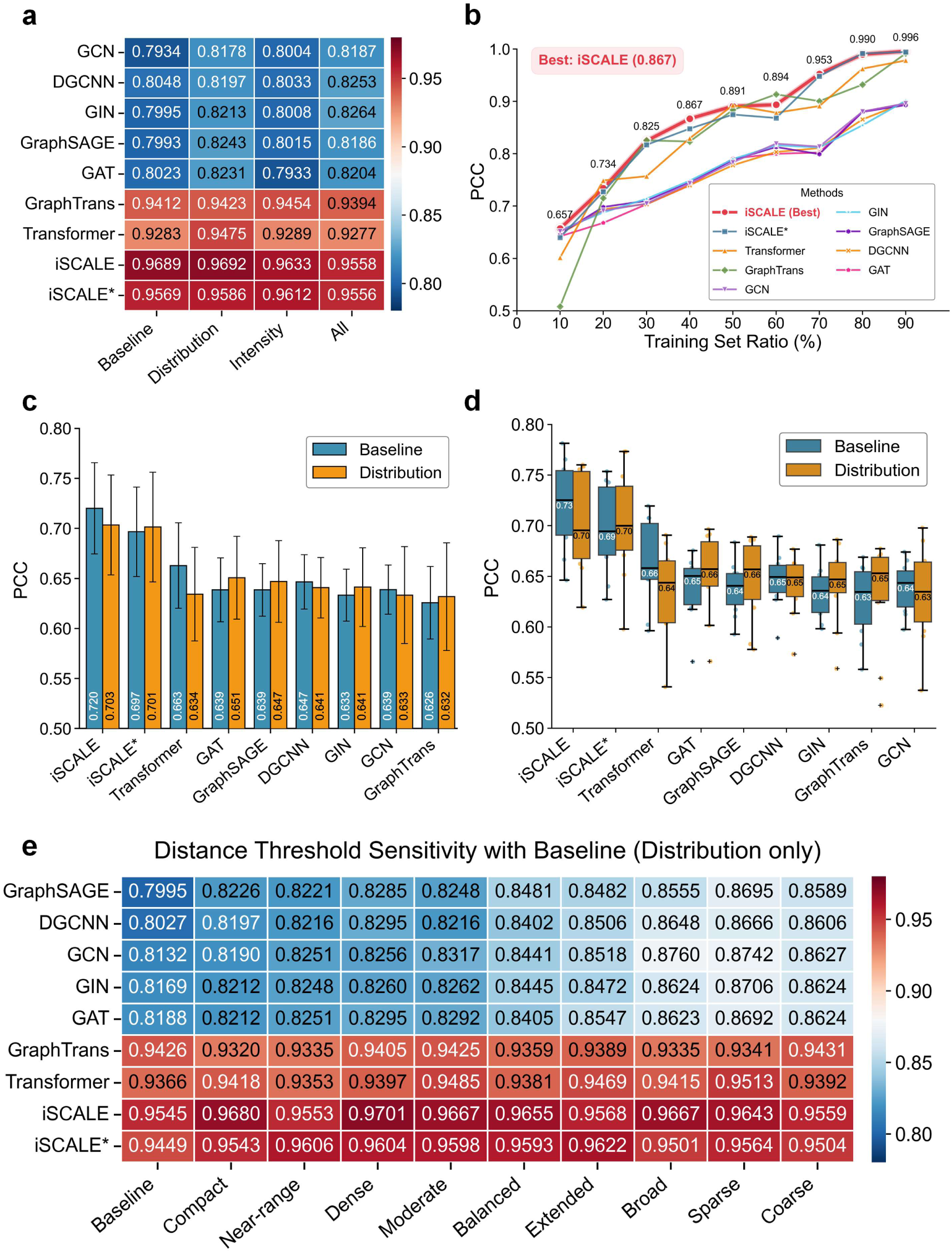
| Benchmark evaluation of protein–RNA binding ΔΔG prediction. Predictions are evaluated by Pearson correlation coefficient (PCC) on a curated benchmark dataset. iSCALE denotes the full model with the auxiliary coupling prediction task; iSCALE* denotes the same architecture without the auxiliary task. Baseline features comprise 20-dimensional one-hot encoding, 21-dimensional PSSM profile, and 1-dimensional conservation score per residue (42 dimensions total). Distribution adds a 7-dimensional multiscale coupling vector per residue, encoding the proportion of RNA nucleotides within each of six distance thresholds. Intensity adds a 1-dimensional coupling intensity scalar. The column of **All** indicates the combination of Baseline, Distribution, and Intensity. (**a**) PCC heatmap across nine existing models and four feature configurations under five-fold cross-validation with random splitting. (**b**) Performance in terms of PCC tested under different training set ratios (10%–90%) for all methods; remaining data are split equally between validation and test sets. (**c**) Mean PCC under PDB-limited splitting, in which at most two mutations per PDB complex are randomly selected for training and remaining samples are split equally between validation and test sets. Results are averaged over 10 independent runs. Bars compare Baseline (blue) and Distribution (orange) features; error bars indicate standard deviation. (**d**) Box plots of the same 10-run PCC distributions as in **c**, showing medians, interquartile ranges, and outliers. Inset values indicate median PCC. (**e**) PCC heatmap for nine distance threshold configurations (columns) across all architectures, using Distribution features only. Each configuration defines six distance thresholds that partition the protein–RNA distance space at different scales (Supplementary Table 1). Configurations are ordered from compact near-range sampling to coarse broad-range sampling.

The training ratio analysis (Fig. 2b) demonstrates iSCALE’s ability to utilize data efficiently. iSCALE achieves a PCC of 0.825 at 30% training data and 0.867 at 40%, while graph neural networks—which use residue contact graphs as their primary structural representation—remain below 0.825 at 70% training data. Across most training ratios, iSCALE outperforms the majority of compared methods, and the performance gap is particularly pronounced at low training fractions (e.g., iSCALE reaches 0.734 at 20%, whereas most graph methods remain below 0.70 at the same ratio). These results suggest that the combination of the state space architecture and structural supervision enables iSCALE to extract generalizable binding-relevant patterns even from limited training data.

We then applied a more stringent protocol, PDB-limited splitting strategy, in which at most two mutations per PDB complex are randomly selected for training, with remaining samples split equally between validation and test sets. Under this protocol, the baseline iSCALE achieves the highest mean PCC (0.720; Fig. 2c) and highest median PCC (∼0.73; Fig. 2d) across 10 independent runs. The addition of distribution features yields a slight decrease for iSCALE (0.703) while improving most other methods, an observation that may reflect the interaction between input-level structural features and block-level structural supervision in the auxiliary task under limited training data. Across all methods, the PDB-limited results show substantially higher variance than the standard cross-validation (Fig. 2d). Despite this, iSCALE maintains the best overall performance and stability among all methods tested.

### 2.3 Sensitivity of prediction performance to distance threshold configuration

The multiscale coupling representation encodes, for each protein residue, the proportion of RNA nucleotides falling within each of six distance thresholds, yielding a 7-dimensional distribution vector (see Methods). To assess how the specific choice of threshold values affects prediction performance, we designed nine configurations that systematically vary the spatial range covered (maximum threshold from 19 to 120 Å) and the sampling density within that range (average inter-threshold spacing from 2.5 to 22 Å; Supplementary Table 1). The configurations span from compact, finely spaced thresholds concentrated in the near range (Compact: 6.5–19 Å) to broadly spaced thresholds extending across the full spatial range (Coarse: 10–120 Å). All architectures are evaluated with Distribution features under five-fold cross-validation (Fig. 2e).

The results reveal three architecture-dependent patterns. First, the five graph-based models show a consistent trend, with performance increasing progressively from compact to coarser configurations. All five achieve their highest PCC at the Sparse (8– 90 Å) or Broad (10–85 Å) configurations, with improvements of 0.050–0.070 over their Baselines. For example, GraphSAGE improves from 0.800 (Baseline) to 0.870 (Sparse), and GCN from 0.813 to 0.876 (Broad). These observations are in line with expectations, as graph-based models construct edges between residue pairs within a fixed distance cutoff (typically 8 Å), so their local message-passing already captures fine-grained spatial relationships. Thresholds in the compact range (6.5–22 Å) overlap substantially with information already encoded in the graph topology, limiting their marginal contribution. Coarser thresholds that extend to 85–90 Å provide complementary information about longer-range spatial organization beyond the reach of local neighborhood aggregation.

The sequence-based methods show a different pattern. iSCALE achieves its highest PCC at the Dense configuration (7–22 Å; PCC = 0.970), iSCALE* peaks at Extended (8–65 Å; PCC = 0.962), and Transformer peaks at Sparse (8–90 Å; PCC = 0.951). Unlike the graph methods, these architectures do not show a monotonic trend toward coarser scales. iSCALE prefers moderate-range thresholds due to its complementary design of both long-range propagation and local-scale supervision. Specifically, iSCALE captures long-range dependencies by propagating information across the full sequence via recurrent state dynamics, while it also learns local spatial information via direct supervision on chunk-level features. In contrast, the Transformer, which lacks sequential inductive bias, benefits from broader spatial context through sparser thresholds.

GraphTrans represents an intermediate case. Its PCC varies between 0.932 and 0.943 across all configurations—the narrowest range (0.011) among all methods—and its Baseline (0.943) is comparable to its best configuration. This relative insensitivity is consistent with the architecture’s combination of local graph message-passing and global attention, which together capture multi-scale spatial information internally, reducing the marginal contribution of external distance features.

Two observations apply across all architectures. First, every method improves over its Baseline under at least one threshold configuration, confirming that distance information at any spatial scale provides useful signal for ΔΔG prediction. Second, the performance gap between the best and worst non-Baseline configurations is modest for the top-performing methods (iSCALE: 0.016; Transformer: 0.016; iSCALE*: 0.017), indicating that once an architecture effectively utilizes spatial information, the specific threshold boundaries matter less than the general principle of encoding distance at multiple scales. The detailed threshold values and design rationale for each configuration are provided in Supplementary Table 1.

### 2.4 Model attention aligns with experimental protein–RNA coupling patterns

To examine whether iSCALE’s learned representations reflect biologically meaningful structural features, we visualized the model’s attention patterns for the Y12Q mutation in the U1A RNA-binding domain from PDB entry 1AUD. The protein mutation exhibits a strong destabilizing effect with a ΔΔG value of 7.17 kcal/mol, serving as a well-characterized case to be observed. Besides, its binding interface with RNA produces prominent coupling intensity peaks along the protein sequence, yielding a distinctive structural signature for evaluating the model’s attention.

We first visualize the two implicit features, spatial coupling distribution and spatial coupling intensity, for the particular case. The multiscale coupling distribution shows how the protein–RNA proximity varies across the sequence. Each residue’s coupling profile encodes the distribution proportion of RNA nucleotides in different distance bins. Three high-contact regions are visible in the heatmap. The green line identifies the mutation site, which nearly coincide with one of the high-contact regions, indicating the rationality of the distribution features. Picking up the nearest nucleotide for each residue, we further calculate the coupling intensity that defined as the inverse of the nearest distance and visualize it in Fig. 3b. The three high-intensity peaks are clearer and correspond to the high-contact regions. Again, the mutation site is close to one of these peaks, suggesting that the mutation effect is closely correlated with the contact distance between mutation site and RNA. In iSCALE we use these two features as both input features and auxiliary supervision signals, so as to encode such structural information into the model.

**Fig. 3.**
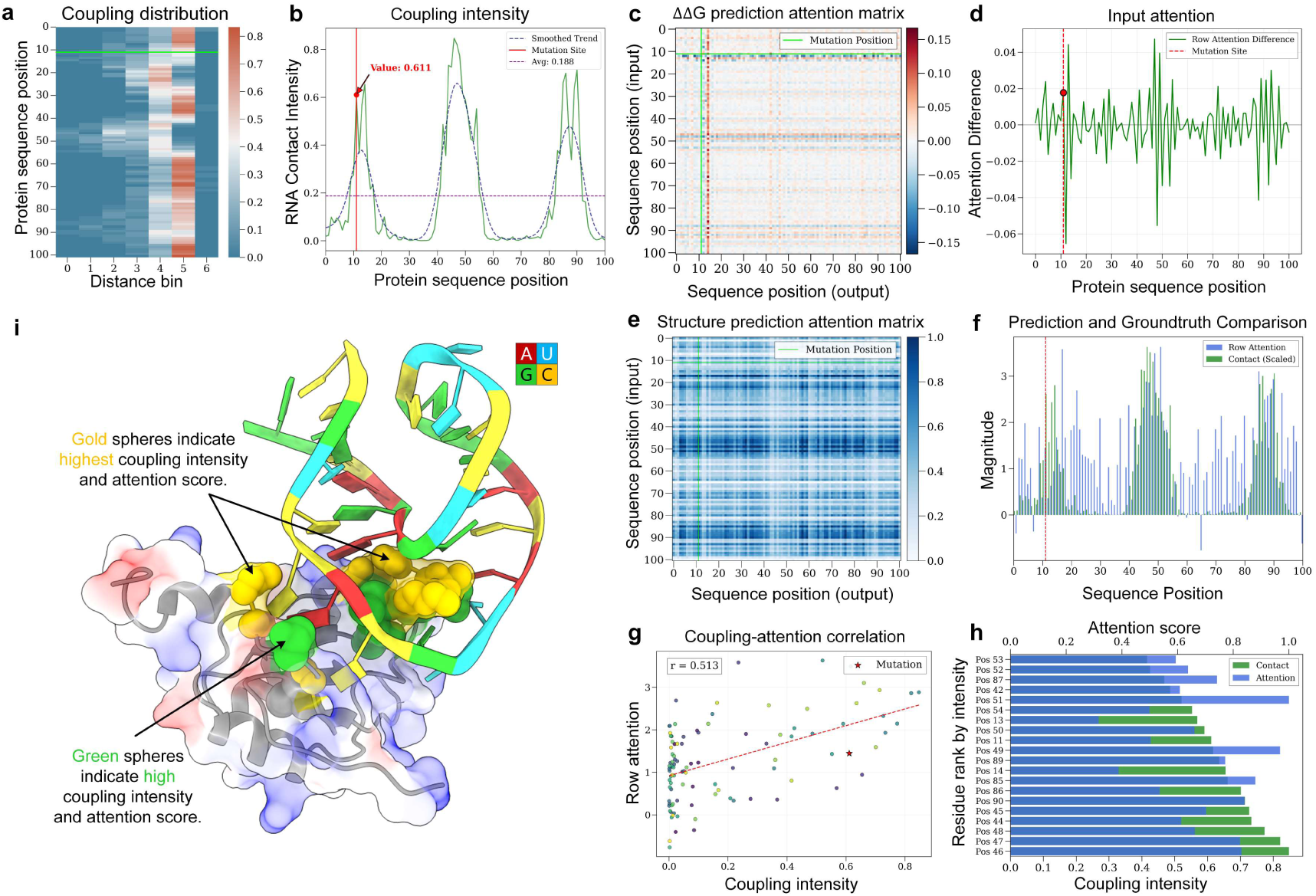
| Correspondence between iSCALE attention patterns and protein-RNA coupling profiles for the Y12Q mutation in the U1A complex (PDB id: 1AUD). Panels are grouped by information source: structural ground truth (a,b), function-oriented attention from the bidirectional concatenation paths (c,d), structure-oriented attention from the isolated wild-type path (e–h), and three-dimensional validation (i). (**a**) Multiscale coupling distribution heatmap. For each protein residue (y-axis), the proportion of RNA nucleotides falling within each of six distance thresholds (8, 10, 15, 20, 30, 50 Å; x-axis bins 0–5) is shown, with bin 6 representing residues beyond 50 Å. Color intensity encodes the proportion (0–1). Green horizontal line: mutation site (Y12). (**b**) Ground-truth RNA coupling intensity along the protein sequence that computed from the wild-type structure. It is inversely proportional to the distance from a residue to its nearest nucleotide. Three prominent peaks correspond to high-contact regions of the binding interface. Red vertical line indicates the mutation site with an intensity value of 0.611, while the average in contrast is only 0.188. (**c**) Attention difference matrix between wild-type and Y12Q mutant sequences, computed from the attention matrices of the bidirectional concatenation paths. Axes represent input (y-axis) and output (x-axis) sequence positions. Green dashed lines: mutation position. A prominent horizontal band at the mutation site indicates that this input position substantially influences all output positions. (**d**) Per-residue attention difference obtained by summing the attention difference matrix along the output dimension. Red dashed line and dot indicate the mutation site and the corresponding value. (**e**) Attention matrix of the wild-type sequence. Green dashed line indicates the mutation position. Three prominent horizontal bands are visible at positions corresponding to the coupling intensity peaks in panel (b). (**f**) Per-residue comparison between row attention from the isolated wild-type path (blue bars) and ground-truth coupling intensity (green bars, scaled to match attention magnitude). Red dashed line indicates the mutation site. (**g**) Scatter plot of ground-truth coupling intensity (x-axis) versus row attention from the isolated wild-type path (y-axis). Each point represents one protein residue; color indicates sequence position. Red dashed line: linear regression. Red star: mutation site. Pearson r = 0.513. **(h)** Top residues ranked by coupling intensity (green) with corresponding attention scores (blue), showing agreement between the two measures at high-contact positions. **(i)** Three-dimensional structure of the protein–RNA complex. Protein surface is colored by electrostatic potential (red: negative; blue: positive); RNA is colored by nucleotide type (A: red; U: yellow; G: green; C: cyan). Gold spheres are the residues with both coupling intensity above the 90th percentile and attention above the 75th percentile. Green spheres are the residues with both measures above the 75th percentile.

We then examined the pairwise interaction scores defined in Section 4.7 (referred to as attention hereafter) to assess whether iSCALE’s learned representations align with the coupling patterns described above. Fig. 3e illustrates the attention matrix of the wild-type sequence. Three prominent horizontal bands can be observed and they align well with the three coupling intensity peaks in Fig. 3b. The attention matrix is then pooled by row to generate per-residue attention magnitude, which is in turn compared against the scaled coupling intensity that used as ground truth. As shown in Fig. 3f and 3g, the two measures show concordant peaks across the sequence, and the correlation across all residues yields a Pearson r of 0.513. In addition, the top-ranked residues by coupling intensity in general corresponds to high attention scores (Fig. 3h). These observations demonstrate that the auxiliary coupling prediction task successfully guides the model to attend to the positions that are in physical proximity to RNA.

The above attention analysis reflects the alignment of model attention to coupling patterns in wild-type structures. However, ΔΔG is a relative value that correlates to the energy change upon mutation rather than the absolute measure of the wild type structure. Therefore, we examined the *attention difference* matrix between wild-type and mutant structures. As shown in Fig. 3c, the attention difference matrix reveals a prominent horizontal band at the Y12Q mutation site, indicating that this input position substantially influences all output positions in the ΔΔG prediction pathway. In addition to the mutation-site band, the matrix shows elevated responses at positions corresponding to the three contact-peak regions identified in Fig. 3b, suggesting that the function-oriented attention pathway has also acquired sensitivity to structurally important positions. The per-residue attention difference (Fig. 3d) confirms this pattern, where the mutation site falls within one of the peak areas. The other peaks also correspond to high-contact regions.

The convergence between the above two attention pathways is notable. The structure-oriented path (Fig. 3e) and the function-oriented path (Fig. 3c) independently identify the same three high-contact regions, despite being computed from different inputs (isolated wild-type versus concatenated wild-type/mutant sequences). This overlap suggests that the auxiliary supervision on the multiscale coupling distribution effectively learns the structural knowledge and propagates to the main ΔΔG prediction branch.

Finally, we mapped the attention scores of protein residues to the protein-RNA structures. The visualization reveals a clear concentration on the high-contact regions of the protein (Fig. 3i). Specifically, residues that score in the top percentiles for both coupling intensity and structure-path attention (gold and green spheres) cluster at the protein–RNA binding interface, rather than being distributed across the protein surface. This spatial co-localization confirms that the model’s high-attention residues correspond to positions that are physically involved in RNA binding.

### 2.5 iSCALE generalizes well to predict protein stability ΔΔG

iSCALE adopts a state space modeling module to learn representations of protein sequences. States are shared across chunks to propagate mutation-induced effects bidirectionally along the sequence so that to model potential long-range dependencies. To investigate whether the state space modeling, as a core design in iSCALE, is a universally effective way for protein learning under mutation scenarios, we assess the performance of iSCALE on the protein stability ΔΔG task. The task relates to only proteins rather than complexes, thus acting an ideal test for assessing solely the effectiveness of the protein representation. Accordingly, the implicit ligand structural features and the auxiliary coupling prediction task in iSCALE are removed. Benchmark datasets are used for evaluations, including S2648 for training and four independent sets for test [24].

We compared iSCALE with nine existing methods specialized for protein stability ΔΔG task (Fig. 4a). At this task, iSCALE achieved the highest PCC on all four benchmarks, reaching 0.831 on S605, 0.839 on S1925, 0.865 on Ssym, and 0.928 on S250. ThermoAGT-GA was the second-best method with PCCs of 0.712, 0.738, 0.848, and 0.907, respectively. The largest gain was observed on S605, which evaluates cross-family generalization, where iSCALE exceeded ThermoAGT-GA by **16.7**%. On S1925, which distributes mutations over four major SCOP structural classes, the improvement is **13.6**%. In addition, iSCALE also achieved the best performance on Ssym with a PCC of 0.865. The Ssym dataset is characterized by the balanced forward and reverse mutation pairs, and it reflects whether a model captures the anti-symmetry of ΔΔG. The good performance of iSCALE on Ssym arises from the bidirectional propagation design in the state space modeling module, so that both forward (wild-type to mutant) and reverse (mutant to wild-type) processes are incorporated. These results suggest that the iSCALE architecture, though originally designed for protein-RNA complexes, generalizes well to the prediction of protein stability ΔΔG. The performance gains are consistently seen across datasets of varying protein families.

**Fig. 4.**
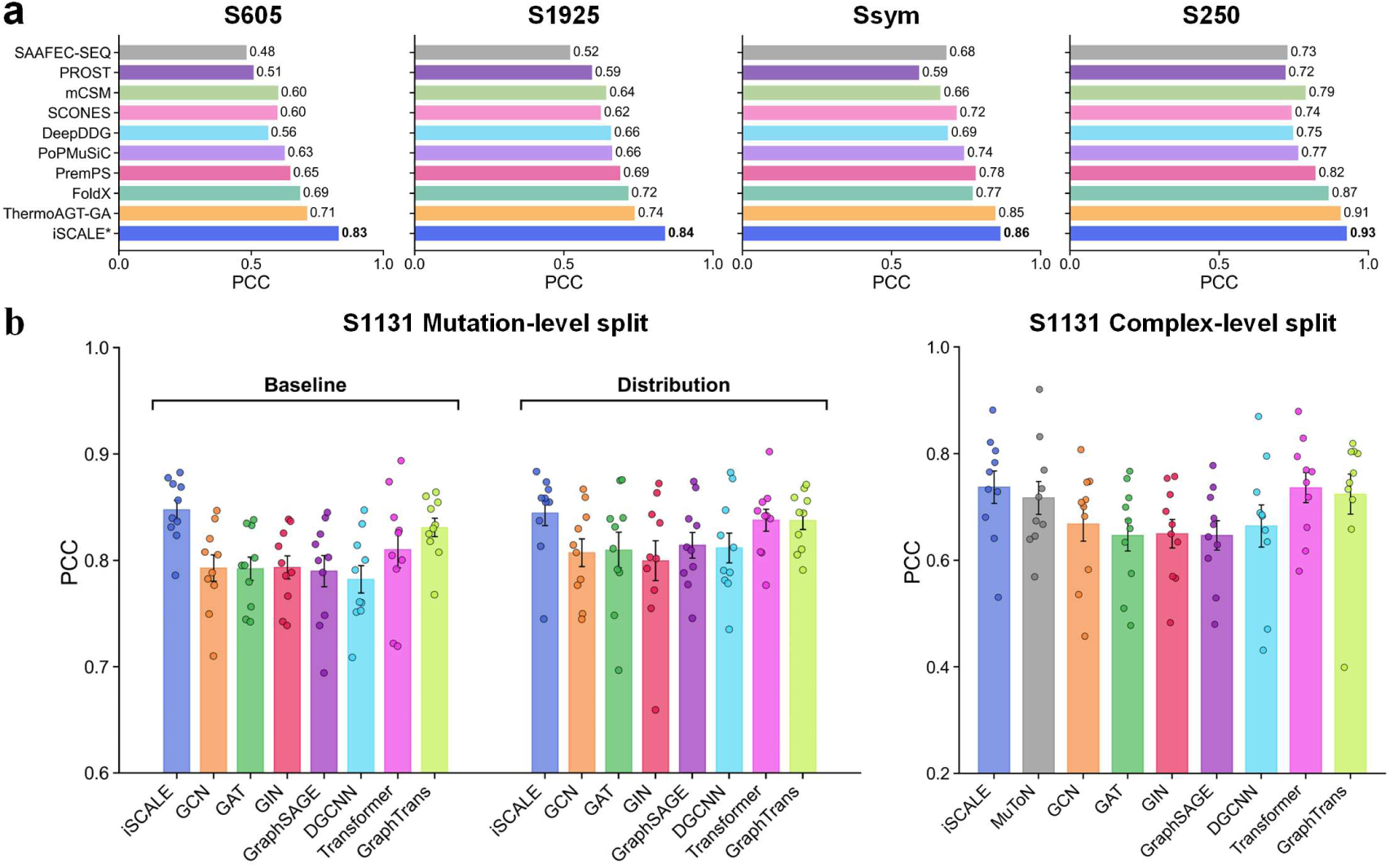
| Performance comparison across different generalization tasks. **(a)** Protein stability ΔΔG prediction task. Horizontal bars show Pearson correlation coefficient (PCC) for iSCALE and nine existing methods across four independent test sets: S605, S1925, Ssym, and S250. All models were trained on the S2648 dataset [24]. **(b)** Protein–protein binding ΔΔG prediction task. The left two charts show model performances on the S1131 dataset of mutation-level random splitting strategy, using baseline and distribution feature configurations, respectively. Colored dots indicate individual values of 10 folds, and error bars indicate standard deviations. The right panel presents model performances on the S1131 dataset of complex-level splitting, in which all mutations from the same PDB complex are assigned to the same fold, ensuring no structural overlap between training and test sets.

### 2.6 iSCALE extends effectively to predict binding ΔΔG in protein-protein complexes

Like protein–RNA interactions, mutations in protein–protein complexes can alter binding affinity and result in the free energy changes (ΔΔG). These changes can be reflected by the interface conformation coupling as well, despite its difference from protein–RNA in physicochemical properties like larger hydrophobic cores and stronger shape complementarity. In this section we assess whether iSCALE indeed captures the inherent structural coupling information and generalizes to broader scenarios. Accordingly, we adapt iSCALE to the PPI task by only replacing the RNA ligand with the partner protein chain, while keeping the model architecture unchanged.

We compared iSCALE against a series of baseline models on the S1131 dataset (“Methods”). Random splitting with ten-fold cross validation is used for an initial regular evaluation. In this regular setting, iSCALE leads with a PCC of 0.8475, approximately 2% above the cutting-edge Graph Transformer [25] and 8.3% above the best graph-based model DGCNN(Fig. 4b). These results suggest that the state space model for modeling mutation effects is not specific to a particular interaction type. Further, to assess whether the multiscale spatial encoding carries relevant information in the PPI context, we incorporated the spatial coupling distribution feature into each baseline model (Fig. 4b, Distribution panel). Performance improvements were observed for all these models including GCN (+1.8%), GAT (+2.2%), GraphSAGE (+3.1%), DGCNN (+3.8%), and Graph Transformer (+3.4%). The consistent improvement that the coupling distribution features bring to other architectures confirms the relevance of multiscale conformational coupling information in PPI. The consistent performance improvements demonstrate the effectiveness of the proposed implicit structural feature.

We then extend the assessment to a more stringent complex-level splitting, in which all mutations from the same PDB complex are assigned to the same fold, ensuring no structural overlap between training and test sets (Fig. 4b, right). This splitting protocol requires the model to predict ΔΔG on entirely unseen protein complexes. Under this protocol, iSCALE still achieves the highest PCC of 0.737, followed by Transformer (0.736). In contrast, MuToN [22], a geometric deep learning method specifically designed for PPI and explicitly modeling binding interface geometry, reaches 0.717. Graph-based models range from 0.647 (GraphSAGE) to 0.669 (GCN). All methods exhibit substantially higher variance under complex-level splitting compared with mutation-level splitting (Fig. 4b, comparing left and right panels), suggesting the difficulty of generalizing across unseen complexes. Nevertheless, iSCALE as a general model without any PPI-specific adaptation perform well on this stringent scenario, outperforming the specialized model MuToN [22] by **2.7**%. These results demonstrate the multiscale conformation coupling better captures the essence of the structural changes than directly modeling geometric structures.

### 2.7 iSCALE discriminates ligand-dependent effects of the same mutation

The same point mutation can produce different functional outcomes depending on the context in which it occurs, such as identical mutations in the same receptor but with different RNA ligands. To investigate whether iSCALE can distinguish such ligand-dependent effects, we analyzed the Y25A substitution in the Hfq RNA chaperone across two related complexes that share the Hfq protein and A7 RNA bound at the distal face, but differ in the presence of an additional RNA ligand. Specifically, we considered the Hfq-A7 binary complex (PDB entry 4HT8 [26]) and the AU6A-Hfq-A7 ternary complex (PDB entry 4HT9 [26]), in which AU6A RNA additionally binds to the proximal face of Hfq, thereby forming a bridged ternary assembly [26]. Despite involving the same amino acid substitution at the same residue position, the Y25A mutation exhibits markedly different effects in the two complexes, with experimental ΔΔG values of 4.18 kcal/mol in the binary complex and -0.28 kcal/mol in the ternary complex. This pair of complexes therefore provides an ideal test case for assessing whether iSCALE can capture mutation-specific patterns that arise from differences in molecular context.

We first assessed the conformational consequences through molecular dynamics simulations. The wild-type binary complex exhibits a compact free energy landscape with a single energy minimum (RMSD ∼0.3–1.1 Å; Fig. 5a), and all three RMSD components stabilize within the first 100–200 ps and remain below ∼4 Å (Fig. 5b). The 4HT9 Y25A mutant produces a similarly compact landscape (RMSD ∼0.3–1.3 Å; Fig. 5e), and the RMSD traces show no sustained drift (Fig. 5f), consistent with the near-neutral ΔΔG. By contrast, the 4HT8 Y25A mutant generates a broader landscape spanning ∼0.5–3 Å in RMSD with multiple dispersed low-energy clusters (Fig. 5g). The RMSD time series reveals pronounced conformational adjustment. As shown in Fig. 5h, the RMSD of the entire complex rises to approximately 6–6.5 Å. The RNA component fluctuates around 2 Å during the first 100–400 ps, increases to about 3.5 Å during 400–700 ps, and then stabilizes near 3 Å during the final 300 ps, while the protein component remains relatively stable at around 2 Å. The larger and more prolonged conformational adjustment in the binary context, compared with the minimal perturbation observed in the ternary context (Fig. 5e,f), is consistent with the substantially higher ΔΔG of 4.18 kcal/mol.

**Fig. 5.**
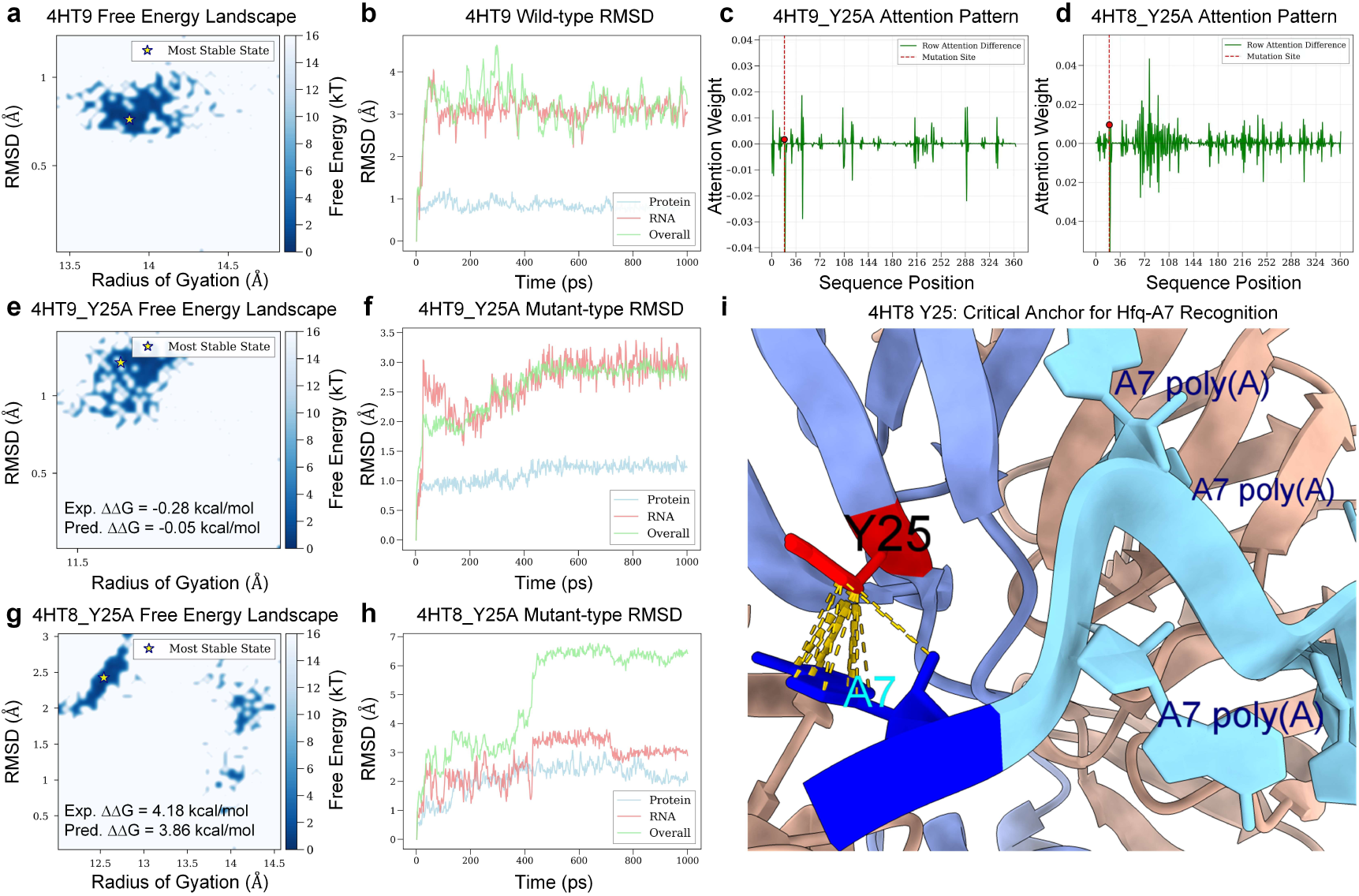
| Context-dependent effects of the Y25A mutation in Hfq across binary and ternary complexes. The same Y25A substitution yields ΔΔG = 4.18 kcal/mol (destabilizing) in the Hfq-A7 binary complex (PDB entry 4HT8 [26]) and ΔΔG = −0.28 kcal/mol (near-neutral) in the AU6A-Hfq-A7 ternary complex (PDB entry 4HT9 [26]). **(a, e, g)** Free energy landscapes of (a) wild-type 4HT9, (e) 4HT9 Y25A, and (g) 4HT8 Y25A from all-atom molecular dynamics simulations, plotted as RMSD versus radius of gyration. Blue regions denote low-energy states, and the star marks the most stable state. **(b, f, h)** RMSD time series of (b) wild-type 4HT9, (f) 4HT9 Y25A, and (h) 4HT8 Y25A. from the same simulations, decomposed into protein (blue), RNA (pink), and overall complex (green). **(c, d)** Per-residue attention difference between wild-type and Y25A sequences in 4HT9, 4HT8 context. Red dashed line and dot indicate the mutation site and its corresponding value. **(i)** Three-dimensional view of the Y25 binding region in the Hfq-A7 complex. Y25 (red) is shown with inter-molecular contacts (gold dashed lines) to neighboring A7 nucleotides (cyan).

Structurally, Y25 stacks with adenosine bases inserted into the R sites on the Hfq distal face [26] (Fig. 5i). In the binary Hfq-A7 complex, the Y25A mutation removes this aromatic stacking interaction at a major RNA-binding site, and the remaining local contacts are insufficient to compensate for its loss. iSCALE successfully captured this strong destabilizing effect by outputting ΔΔG as 3.86 kcal/mol, which is close to the ground truth ΔΔG 4.18 kcal/mol measured in experiments. In the ternary AU6A-Hfq-A7 complex, however, AU6A engages the spatially distinct proximal face through an additional interaction network that is largely unaffected by Y25A [26]. These preserved interactions may buffer the energetic penalty associated with perturbation of the distal Y25-adenosine contacts, resulting in a near-neutral net ΔΔG of −0.28 kcal/mol in experiments. In this case, iSCALE predicted ΔΔG as −0.05 kcal/mol, showing its strong ability in discriminating ligand-dependent effects of the same mutation.

The structure-guided auxiliary task contributes directly to prediction accuracy in this context-dependent case. With the multiscale coupling auxiliary task, iSCALE predicts the 4HT8 Y25A ΔΔG as 3.86 kcal/mol, close to the experimental value of 4.18 kcal/mol (error: 0.32 kcal/mol). In contrast, removing the auxiliary task results in the prediction dropping to 3.11 kcal/mol (error: 1.07 kcal/mol). These results suggest that the structural supervision through the coupling prediction task enables the model to capture residue-level interaction patterns underlying context-dependent effects, which are difficult to infer from ΔΔG labels alone.

In addition to the accurately predicted ΔΔG, iSCALE exhibits attention patterns that support the predictions. For the ternary complex (Fig. 5c), the model attended to a small portion of residues and most positions were considered to cause minor impacts on the binding affinity. In contrast, the attention difference in the binary complex (Fig. 5d) is more pronounced across residues, suggesting that the binary one is more prone to mutations in most positions.

### 2.8 iSCALE distinguishes distinct mutation-specific destabilizing effects within the same complex

A protein-RNA complex can exhibit different destabilizing effects depending on the specific mutation sites. To examine whether iSCALE is capable of distinguishing destabilizing effects of different mutations, we selected three single-site mutants of the RNA-binding protein U1A (PDB entry 1AUD) for detailed analysis. As shown in the first row in Fig. 6, the three substitution mutations, namely Y12Q, G52A, and F55L, are all located within high-contact-intensity regions of the protein–RNA interface. For each mutation, we performed the attention pattern analysis and molecular dynamics (MD) simulations.

**Fig. 6.**
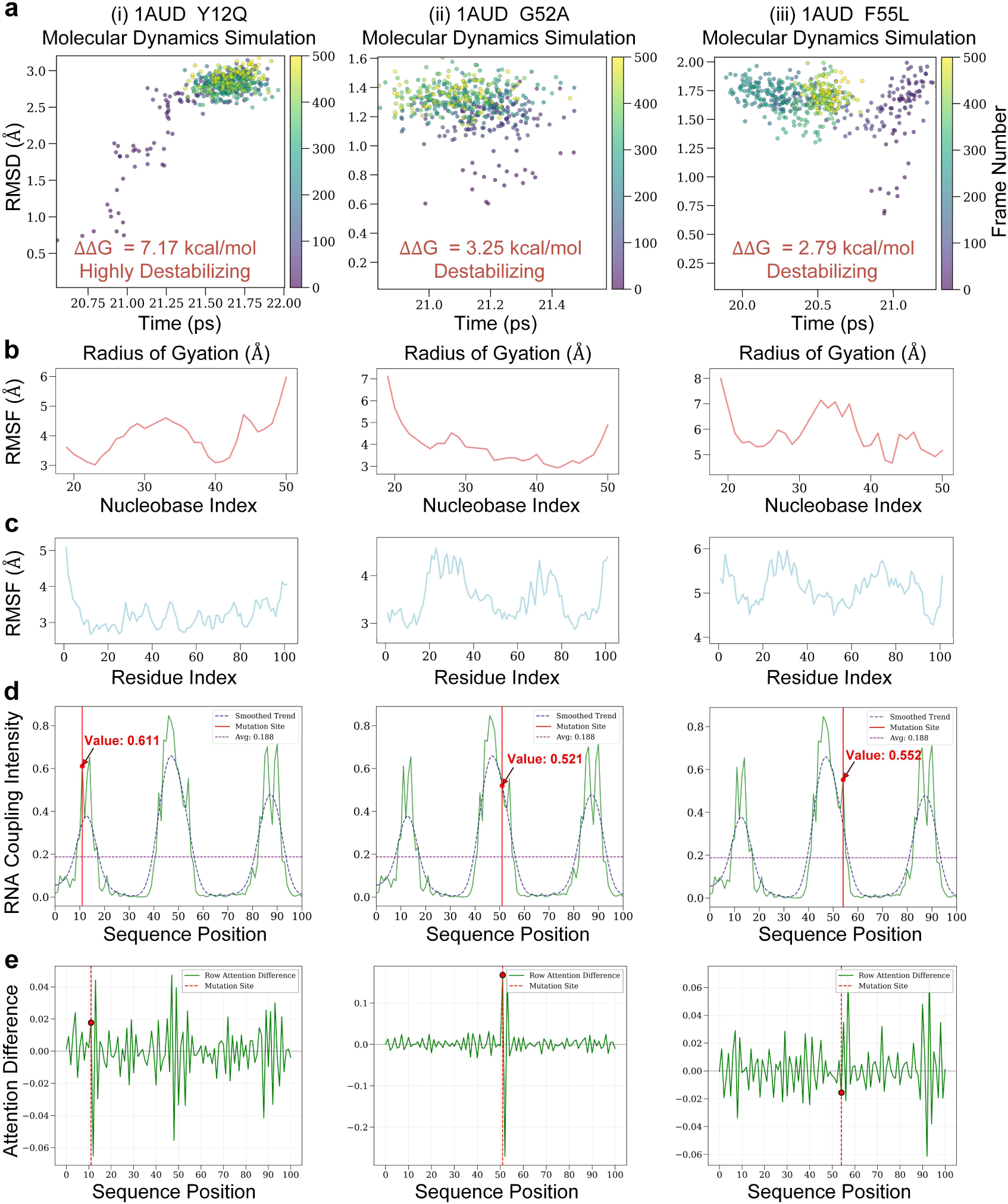
| Molecular dynamics simulation and attention analysis of three destabilizing mutations in the U1A–RNA complex (PDB entry 1AUD). Three mutations at high-coupling-intensity regions—Y12Q (i), G52A (ii), and F55L (iii)—are analyzed across five dimensions. Experimental ΔΔG values are 7.17, 3.25, and 2.79 kcal/mol, respectively. **(a)** Conformational trajectories from molecular dynamics simulations, plotted as RMSD versus radius of gyration, with colors indicating the chronological order of frames. **(b, c)** Root mean square fluctuation (RMSF) of RNA nucleotides (b) and protein residues (c) extracted from the same trajectories, quantifying per-residue flexibility upon mutation. **(d)** Ground-truth RNA coupling intensity profile along the protein sequence, computed from the wild-type crystal structure. Red vertical lines mark the mutation sites. The dashed purple line indicates the sequence-averaged intensity. **(e)** Attention patterns for different mutations, calculated as the model attention difference between wild-type and mutant sequences. Regions with large absolute values indicate positions where the model attention is substantially altered in mutant sequence compared with the wild-type one.

First, the mutation Y12Q is a highly severe destabilizing variant with a ΔΔG of 7.17 kcal/mol. In this case, iSCALE’s attention pattern the attention difference spans multiple high-contact regions along the sequence, rather than concentrating solely at the mutation site (Fig. 6a, Row V). Structurally, Y12 participates in planar stacking interactions with RNA bases at the binding interface [27], and its replacement with glutamine would be expected to eliminate this aromatic stacking contribution, consistent with the high ΔΔG value. MD simulations generate a conformational trajectory covering a broad RMSD range (∼0.5–3 Å), with the early frames tracing an extended path before gradually clustering (Fig. 6a, Row I). The corresponding RMSF profiles reveal elevated flexibility across both protein and RNA chains (Fig. 6a, Rows II–III). The co-occurrence of distributed attention changes at high-contact positions, broad conformational adjustment in the trajectory, and elevated flexibility across both chains suggests that this severe mutation perturbs the binding interface at multiple sites, and the model allocates attention to positions where structural rearrangement is substantial.

By contrast, for the mutation G52A (ΔΔG = 3.25 kcal/mol), its attention pattern is concentrated near the mutation site, with minimal changes elsewhere (Fig. 6b, Row V). G52 is located in the loop 3 region, which participates in key interactions at the protein– RNA interface [27]. The G→A substitution introduces a side-chain methyl group that restricts the backbone conformational freedom characteristic of glycine, and the localized attention is consistent with a perturbation that remains spatially confined. The trajectory remains within a narrow RMSD range throughout the simulation (Fig. 6b, Row I), and the RMSF profiles stay relatively uniform (Fig. 6b, Rows II–III), confirming that both the model’s attention and the simulated structural response point to a localized effect near the mutation site.

For F55L (ΔΔG = 2.79 kcal/mol), the attention difference shows moderate changes distributed across the sequence, with multiple regions exhibiting altered attention rather than a single dominant peak (Fig. 6c, Row V). F55 participates in intermolecular stacking interactions with RNA base A44 at the binding interface [27], and the F→L substitution replaces the aromatic ring with a branched aliphatic side chain, which would be expected to disrupt these stacking contacts. The trajectory reveals two distinct conformational clusters with a clear transition between them (Fig. 6c, Row I), and the RMSF profiles show region-specific fluctuation changes in both chains (Fig. 6c, Rows II–III). The two-state conformational behavior, region-specific flexibility changes, and distributed attention changes together suggest that this disruption propagates beyond the immediate mutation site, involving coordinated rearrangement at multiple positions along the interface.

A notable observation arises from comparing G52A and F55L. Both mutations are located at the central contact intensity peak and are spatially proximate (Fig. 6, Row IV), yet they produce clearly distinct attention profiles and conformational behaviors (Fig. 6, Rows I, V). G52A shows localized attention and rapid structural convergence, while F55L shows distributed attention and a two-state trajectory. This contrast indicates that the model’s attention differences are sensitive to the specific identity and functional consequence of each mutation, rather than simply reflecting proximity to the binding interface. Together, these three cases illustrate that iSCALE’s learned representations carry mutation-specific information that goes beyond the structural features explicitly provided as input.

## III. Discussion

Quantifying mutation-induced changes in protein-RNA binding ΔΔG provides basis for understanding pathogenic outcomes. Traditional experimental measurements are impractical to exhaustively quantify numerous variants, which motivates the development of efficient data-driven methods. Existing methods have adopted sequence-based and structure-based methods. However, sequence-based models fall short in capturing subtle mutation-induced signals and are not capable of incorporating mutation-induced structural changes. In contrast, structure-based models, though characterizing local structural environments more faithfully, may overfit to local structural noise rather than learning a general geometric pattern.

Motivated by the above drawbacks, we introduced iSCALE, an interpretable and generalizable deep learning model for the prediction of mutation-induced binding affinity changes. The key conceptualization is to encode binding-partner structures implicitly in residue features, rather than modeling it explicitly with conventional geometric learning methods. A state space modeling module is adopted to propagate mutation effects across long protein sequences. This design models the long-range dependencies efficiently while injecting structural information into residues without overfitting to excessively detailed geometric patterns. These together provide a biologically motivated way to connect local interface geometry with global sequence-level reasoning.

iSCALE consistently surpassed eight existing methods on the benchmark protein-RNA binding ΔΔG prediction dataset, achieving a high PCC score of 0.969. In addition, iSCALE remained robust when evaluated on reduced training data, indicating a high data utilization efficiency upon training. Likewise, the model’s consistently superior performance under the more stringent PDB-limited splitting strategy suggests that iSCALE is capable of learning inherent binding patterns rather than memorizing protein-dependent shortcuts. Of note, adding the multiscale coupling distribution feature improved performances of almost all models, confirming the universal applicability of the proposed module across models.

A particular important finding is that iSCALE remains discriminative in some difficult scenarios, where the same mutation produces different outcomes depending on the ligand, or mutations occurred within the same complex but corresponding to different destabilizing effects. In the Hfq system, the identical Y25A mutation showed markedly different effects in the binary and ternary complexes, and iSCALE captured this distinction by producing different attention patterns. Similarly, in the U1A complex, the model distinguished multiple destabilizing mutations at different sites, even when they were spatially close. These particular cases indicate that iSCALE does not learn a trivial signal that merely approximates the interface structures. Instead, it recognizes mutation-specific structural patterns and context-dependent conformational coupling. This discrimination is essential for practical applications, where the biological effect of a mutation is often determined by the full molecular environment, not by residue identity alone.

iSCALE exhibits strong generalization to mutation-related tasks other than protein-RNA complexes, suggesting that the learned representations capture essential patterns of mutation-associated changes. These changes are not restricted to a single biomolecular setting. When adapted to protein stability ΔΔG prediction, the model achieved consistently superior performances to existing specialized methods across multiple benchmark sets. This observation underlines the effectiveness of the protein representation learned by the state space module alone. Likewise, when transferred to protein-protein binding ΔΔG prediction, i.e., the ligand becomes a protein rather than an RNA, iSCALE maintained strong predictive accuracy and remained robust under stricter complex-level splitting. These results show that iSCALE offers a unified approach for predicting stability or binding affinity changes upon mutations. Despite the apparent differences among these tasks, iSCALE adopts a uniform modeling strategy featured by the multiscale spatial encoding, requiring no task-specific adjustments. The model’s consistency across tasks is in line with the shared biological principles underlying diverse molecular interactions.

Despite the above advantages, iSCALE faces several potential limitations. First, the current framework relies on available structural information to generate the multiscale spatial encoding, thus cannot being directly applied to cases without known structures. Whether virtual structures by AlphaFold-3 or other tools are equally effective warrant further investigations. Second, the present analysis focuses on single-site mutations, whereas real biological systems often involve combinatorial variants and coupled mutational effects. Finally, although the model generalizes well across related tasks, prospective validation on larger protein–RNA datasets, as well as more diverse ligand types like drugs and peptides, will be necessary to assess its robustness in broader experimental settings.

## IV. Methods

### 4.1 Datasets

We evaluated iSCALE on multiple benchmark datasets covering three mutation-related tasks, including protein-RNA binding ΔΔG prediction, protein stability ΔΔG prediction, and protein-protein binding ΔΔG prediction. For protein-RNA binding ΔΔG prediction, we adopted the S788 dataset from PRITrans [13], which contains 788 experimentally validated missense mutations covering 78 protein-RNA complexes. The dataset was constructed from 394 original mutations and their thermodynamically derived reverse counterparts, with ΔΔG values ranging from −7.17 to +7.17 kcal/mol. For protein stability prediction, we used the S2648 dataset as the training set. It contains 2,648 non-redundant single-point mutations from 131 globular proteins derived from the ProTherm database [28]. Four independent test sets were used for evaluation [24]: S605 (605 mutations from 58 proteins), S1925 (1,925 mutations from 55 proteins distributed over four major SCOP structural classes), S250 (250 forward and reverse mutations from 9 proteins with both wild-type and mutant structures available), and Ssym (684 mutations with balanced forward and reverse pairs). For protein-protein binding affinity prediction, we adopted the S1131 dataset [22]. It is a subset of SKEMPI v2.0 [29] and contains 1,131 single-point mutations at protein-protein binding interfaces.

We curated all the above datasets with a unified pipeline. For each sample, the corresponding wild-type three-dimensional structure was retrieved from the RCSB Protein Data Bank [30]. Since experimentally resolved structures frequently contain atomic clashes and missing residues, we further employed FoldX 5.0 [6] to repair the wild-type structures using the RepairPDB command. The mutant structures were then generated via BuildModel. For the protein features, position-specific scoring matrices (PSSMs) were generated using PSI-BLAST (BLAST+ 2.16.0) [31] against the SwissProt database with three iterations, and residue-level conservation scores were derived from the resulting PSSM profiles. For protein-RNA and protein-protein complexes, protein and ligand chains were separated to obtain the protein-centric representations.

To evaluate the generalizability of models, we adopted various splitting strategies depending on the task. For the protein-RNA binding task (S394), five-fold cross-validation with random splitting was used as the primary benchmark. To further assess performance under data-scarce conditions, a PDB-limited splitting strategy was employed. Specifically, for each PDB complex, at most two mutations were randomly selected for training, and all remaining samples were split into validation and test sets at a 1:1 ratio. This procedure was repeated across 10 independent runs with different random seeds to reflect general estimates. For the protein-protein binding task (S1131), ten-fold cross-validation with random splitting served as the regular evaluation. A more stringent PDB-based splitting was additionally applied, in which samples were grouped by PDB identifier and entire groups were assigned to folds such that all mutations from the same complex remained within a single fold, ensuring no structural overlap between training and test sets. For the protein stability task, models were trained on S2648 and evaluated on four independent test sets (S605, S1925, S250, and Ssym).

### 4.2 Problem formulation

Given a wild-type protein sequence *P*_wt_, a mutant sequence *P*_mt_ differing from *P*_wt_ by a single amino acid substitution, and the three-dimensional structure of the complex containing the protein and its binding partner (RNA or protein), the task is to predict the mutation-induced change in binding free energy:

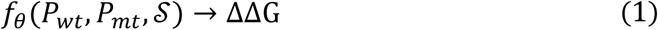

where *S* denotes the atomic coordinates of the complex structure, from which both protein residue positions and protein–ligand distances are derived, and *ΔΔG* = *ΔG*_mt_ − *ΔG*_wt_ is the difference in binding free energy between the mutant and wild-type complexes.

### 4.3 Feature construction and graph representation

#### 4.3.1 Node feature construction

We convert the protein sequence *P* = {*a*_1_, *a*_2_, …, *a*_LP_} into feature representation **X** = {**x**_1_, **x**_2_, …, **x***_LP_*}, where each amino acid residue *a_i_* corresponds to a feature vector:

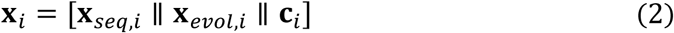

where **x***_seq,i_* ∈ ℝ^20^ denotes the one-hot encoding of amino acid type, and **x***_evol,i_* ∈ ℝ^21^ is an evolutionary feature vector that contains 20-dimension PSSM features and 1-dimension conservation score. ***c****_i_* represents multiscale coupling features that encodes spatial interaction information between protein and ligand. The three components correspond to the direct information of sequences, the indirect information of evolution, and the implicit information of structures, respectively, providing comprehensive and complementary feature sources for the state space model.

#### 4.3.2 Protein-centric graph construction

Considering the different types of binding ligands, we construct a protein-centric residue graph where only the protein is explicitly modeled as a graph and the ligand is implicitly encoded into the graph node features. Specifically, the protein-centric graph *g_P_* = (*v_P_, Ɛ_P_*) is constructed:

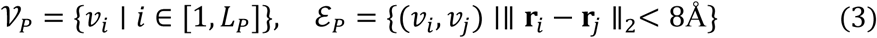

where **r***_i_* is the *C*α atom coordinate of residue *i*. A pair of residues with a distance less than 8Å are connected by placing an edge. For edge features, we employ sinusoidal positional encodings **e***_ij_* = PE(*d_ij_*), where *d_ij_* =ǁ **r***_i_* − **r***_j_* ǁ_2_ is the inter-residue distance. For the binding ligands (RNA or protein), we only utilize their atomic coordinate information to calculate protein-ligand coupling without explicitly modeling the ligand structure. This is achieved by the aforementioned multiscale coupling features **c***_i_* . This protein-centric representation reduces the complex protein-RNA bipartite graph problem to a one-sided protein graph problem while preserving RNA interaction information through conformational coupling features.

### 4.4 Multiscale spatial encoding module

The multiscale spatial encoding module implicitly encodes the ligand structural information into protein representations. It takes forms of either a scalar value representing spatial coupling intensity or a distribution vector indicating the spatial coupling distribution.

#### 4.4.1 Spatial coupling intensity

Node-level coupling intensity is based on a sigmoid function of minimum distance:

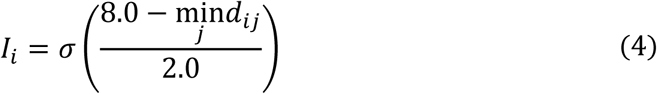

where 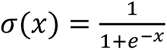 is the sigmoid function. This S-shaped function converts distance to coupling intensity, which aims to reflect the physical characteristics of molecular interaction strength decay with distance.

#### 4.4.2 Spatial coupling distribution

For a protein residue *i*, its multiscale coupling feature vector is defined as:

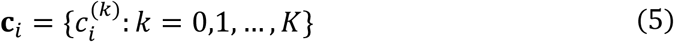

where the *k*-th component is:

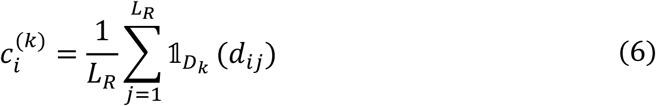

Here *d_ij_* is the distance between protein residue *i* and RNA nucleotide *j*, *L_R_* is the RNA sequence length, and 1*_D_k__*(⋅) is the indicator function for distance interval *D_k_*:

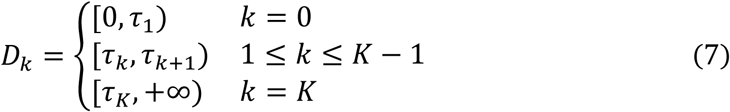

where **τ** = {8Å, 10Å, 15Å, 20Å, 30Å, 50Å} is the distance threshold set. The residue-level coupling distribution is grouped to form a chunk-level distribution when used as supervisory labels for auxiliary tasks. This prevents model from overfitting to particular residue-level structural details.

### 4.5 Bidirectional state space model

We chose state space models as the technical foundation because their *A*-matrix exponential decay properties naturally align with the physical characteristics of conformational propagation. Each SSD layer applies a linear projection, a causal depthwise convolution (kernel size 4), and a SiLU nonlinearity to produce an input signal **x** ∈ ℝ*^L×P^*, two encoding vectors **B**, **C** ∈ ℝ*^L×N^*, a timescale parameter **Δ** ∈ ℝ*^L^*_>0_, a gating vector **z**, and a per-head skip parameter *D*, where *L* is the sequence length, *P* the head dimension, and *N* the state dimension. A per-head learnable rate *A* < 0 controls exponential decay. **B** and **C** are shared across all heads. The core recurrence of the state space model is:

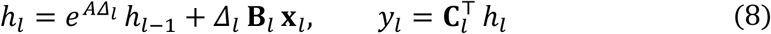

where *ℎ_l_* is a state vector carrying a compressed summary of all inputs up to position *l*, updated at each step by decaying the previous state (governed by *A* and *Δ_l_*) and incorporating the new input (weighted by **B***_l_*). The output *y_l_* is read out by projecting the state through **C***_l_* . This provides a mathematical framework consistent with the distance-dependent decay of conformational coupling influence.

For efficiency, this sequential recurrence is decomposed into within-chunk pairwise interactions and cross-chunk state propagation [23]. The sequence is divided into non-overlapping chunks of *Q* residues. Within each chunk, the recurrence can be unrolled into a closed-form weighted sum:

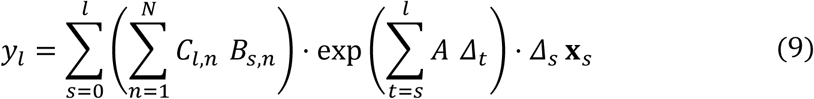

where *l* and *s* index positions within a chunk (from 0 to *Q* − 1). The pairwise score ∑*_n_ C_l,n_ B_s,n_* quantifies the interaction between positions *l* and *s* in the *N*-dimensional encoding space. The causal decay exp(∑*AΔ*_t_), with *A* < 0, ensures that more distant positions contribute exponentially less, modulated by the data-dependent timescale *Δ*. This pairwise form yields a *Q* × *Q* interaction structure analogous to the query-key product in self-attention.

Equation (9) captures interactions only within the same chunk. To propagate information across chunk boundaries, the state vector ℎ at each boundary carries a compressed summary of all earlier inputs, accumulated using the same recurrence in equation (8). Each position *l* then receives an additional contribution by projecting this state through **C**_l_, scaled by cumulative decay. This is added to *y*_l_ from Equation (9), and the combined output is modulated by the gating vector: output = (*y* + *D* ⋅ **x**) ⊙ SiLU(**z**).

The bidirectional differential mechanism captures the directionality of conformational changes through:

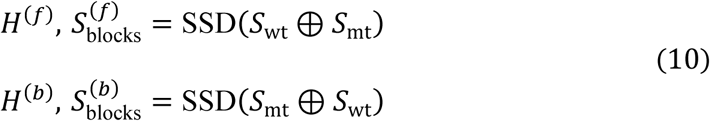

where ⊕ represents sequence concatenation, and superscripts (*f*) and (*b*) denote forward and reverse paths, respectively. In each concatenation, positions in the second segment can attend to all positions in the first segment through the pairwise interactions and state propagation described above, while positions in the first segment are encoded largely independently. The differential conformational coupling effect is calculated as:

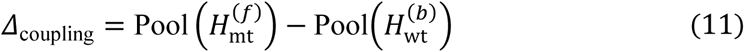

Each SSD sublayer further consists of a forward and a backward pass with separate parameters, whose outputs are combined through a learned gate **g**:

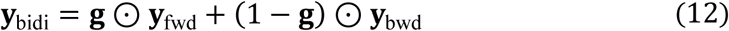

where **g** is computed from the concatenation of **y**_fwd_ and **y**_bwd_ . This bidirectional processing is orthogonal to the sequence concatenation above: the concatenation controls which variant is encoded in whose context, while the bidirectional passes ensure each sequence is read from both directions.

The SSD computation produces intermediate block-level states as a byproduct of the cross-chunk recurrence: the state ℎ at each chunk boundary constitutes a fixed-dimensional summary of all residue-level inputs within and before that chunk. For chunk *b* from the wild-type path, a representation vector *s_b_* ∈ ℝ^*d*_inner_^ is obtained by averaging the states along the head dimension:

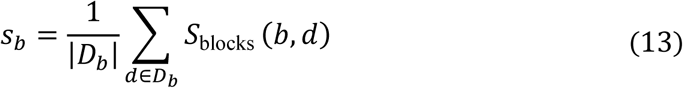

where *D_b_* is the set of head indices and *S*_blocks_(*b*, *d*) is the state of chunk *b* at head *d*. Each *s_b_* is then passed through two separate MLP heads:

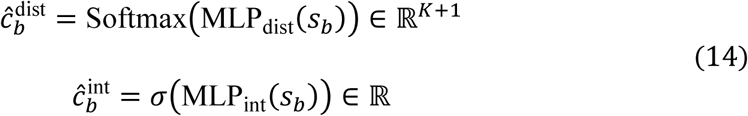

where 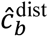 is the predicted coupling distribution over *K* + 1 distance intervals defined by the threshold set τ (Section 4.4), 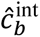 is the predicted coupling intensity, and *σ* denotes the sigmoid function.

### 4.6 Dual-task prediction module

iSCALE predicts *ΔΔG* as the main task, while it also predicts the chunk-level spatial coupling distribution as an auxiliary task. iSCALE contains three paths at each SSD layer, including a forward concatenation [**S***_wt_*; **S***_mt_*], a backward concatenation [**S***_mt_*; **S***_wt_*], and a wild-type sequence *S_wt_* alone. The first two paths produce the differential representation for the main *ΔΔG* prediction. The third path is dedicated to the auxiliary coupling prediction task. The parameters are shared across all three paths to ensure that the block-level states used for auxiliary supervision are produced by the same learned dynamics as those driving the main prediction, so that the structural knowledge captured by the auxiliary task directly shapes the representations used for *ΔΔG* inference.

iSCALE is trained end-to-end with joint optimization of main and auxiliary tasks. The total loss function is defined as a weighted combination of main and auxiliary losses:

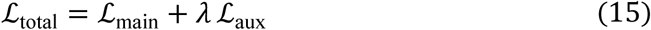

The main task uses mean squared error loss: 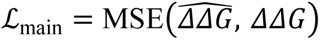. *λ* denotes the weight of auxiliary tasks and was set to 0.2 in this study. Auxiliary task loss includes joint loss of block-level coupling distribution prediction and coupling coupling intensity prediction. It aggregates predictions from all SSD layers:

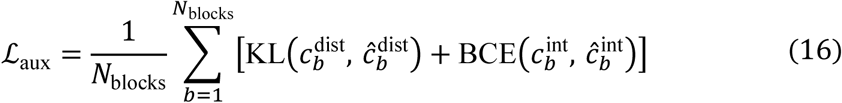

where 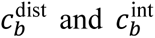 are the ground-truth block-level coupling distributions and intensities computed from three-dimensional coordinates, and 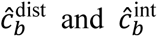 are the corresponding predictions at chunk *b*.

### 4.7 Pairwise interaction score

The pairwise score 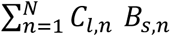 in Equation (9) quantifies the data-dependent interaction between output position *l* and input position *s* within each chunk. To characterize this interaction structure across the full sequence, we extract the encoding vectors **B**, **C** ∈ ℝ*^L×N^* from the first SSD layer’s forward sublayer and define

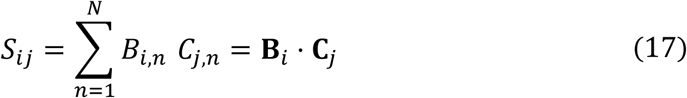

which isolates the data-dependent encoding alignment between positions *i* and *j*, computed over the full sequence length without the causal decay 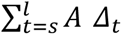 or timescale modulation *Δ_s_* applied within each chunk. The per-residue magnitude *ā_i_* = 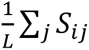 measures how strongly position *i*’s encoding aligns with the global encoding direction across all positions. To identify where a mutation alters this encoding structure, we compute the score difference Δ*S* = *S_mt_* − *S_wt_* between the mutant and wild-type segments extracted from the differential paths in equation (10). In the Results (Sections 2.4, 2.7, and 2.8), we refer to *S* and its derived quantities as *attention* for brevity, reflecting the structural correspondence between the pairwise score in Equation (9) and the query-key product in Transformer self-attention.

### 4.8 Training details

The model was trained on an NVIDIA GeForce RTX 4090 GPU using the PyTorch framework. The Adam optimizer was used with a learning rate of 8 × 10^−4^ and a weight decay of 1 × 10^−5^. A plateau learning rate scheduler was used with a decay factor of 0.8 and a patience setting of 15 epochs. The batch size for training was set to 16. The maximum training epoch was 300, and early stopping was employed with a patience of 40 to prevent overfitting. Other parameter settings regarding network architecture include a hidden dimension 64, three SSD layers, a state dimension of 32, a convolution kernel size of 4, a chunk size of 32, and a dropout rate of 0.1.

### 4.9 Evaluation metrics

In this study, four metrics are used to assess the model performance, including PCC, RMSE, MAE, and *r*_att_cont_. PCC measures the linear correlation between the predicted and experimental ΔΔG values:

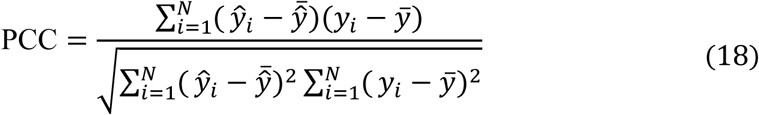

In this study we also use PCC to evaluate the correlation between the learned attention scores and coupling intensity, so that to reflect the model’s understanding of conformation coupling.

RMSE quantifies the average magnitude of the prediction error by taking the square root of the average squared differences between predicted and observed values, while MAE represents the average of the absolute differences between them.

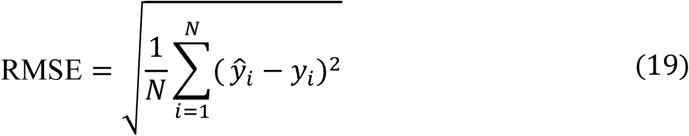

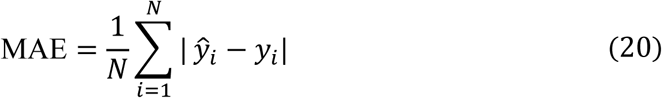

## Data Availability

All data supporting the findings of this study are available in the article, supplementary materials, and source data files. The protein-RNA interaction dataset S394 was obtained from PRITrans [https://github.com/cuifengLI/PRITrans]. The protein thermostability datasets S2648, S605, S1925, S250, and Ssym were downloaded from ThermoAGT [https://github.com/mahan-fcb/ThermoAGT]. The protein-protein binding affinity dataset S1131 was obtained from MuToN [https://github.com/zpliulab/MuToN]. Wild-type protein structures were retrieved from the RCSB Protein Data Bank [https://www.rcsb.org]. Structure repair and mutant structure generation were performed using FoldX 5.0 [http://foldxsuite.crg.eu/]. Position-specific scoring matrices were generated using PSI-BLAST (BLAST+ 2.16.0) [https://blast.ncbi.nlm.nih.gov/Blast.cgi] against the SwissProt database [https://www.uniprot.org]. The processed protein-RNA dataset and source data are available via Zenodo at https://doi.org/10.5281/zenodo.21972360.

## Code Availability

The source codes of iSCALE, including data preprocessing, model training, and evaluation scripts are available on GitHub at https://github.com/ssmiter/DualPRI. The code repository includes detailed usage instructions, example data, and scripts for reproducing experimental results.

## Acknowledgements

This work was supported by the National Natural Science Foundation of China (NO. 62306293), the Natural Science Foundation of Shandong Province (NO. ZR2025MS1069), the Fundamental Research Funds for the Central Universities (NO. 202641011), the Youth Innovation Technology Project of Higher School in Shandong Province (NO. 2025KJH006), and the Qingdao Key Technology Breakthrough Project (NO. 26-1-1-gjgg-66-nsh).

