## Supplementary Information for "Predicting Protein-RNA Binding Affinity Changes via Spatial Coupling-Aware State Space Modeling"

### SUPPLEMENTARY MATERIAL

#### Supplementary Note 1: Distance threshold configuration design

The multiscale spatial encoding module requires defining a set of distance thresholds that partition the protein–ligand distance space. Two design parameters determine the representation: the number of thresholds and the specific threshold values.

*Number of thresholds.* We fixed the number of thresholds at six throughout all experiments, yielding a 7-dimensional distribution vector per residue (Section 4.4). This choice balances representational capacity against overfitting risk: too few thresholds collapse the multiscale structure into a coarse approximation, while too many approach a continuous distance representation that is computationally expensive and difficult to aggregate at the chunk level required by the auxiliary task (chunk size $Q = 32$ residues). Six thresholds provide sufficient granularity to distinguish near-range, intermediate, and long-range spatial relationships while remaining tractable for chunk-level supervision.

*Threshold value selection.* To assess whether specific threshold values substantially affect prediction performance, we designed nine configurations that systematically explore the space of possible threshold sets (Supplementary Table 1). We varied two key properties: (1) the spatial range covered, defined by the minimum and maximum thresholds, and (2) the sampling density, defined by the average spacing between consecutive thresholds. The nine configurations span from compact near-range sampling (Compact: 6.5–19 Å, average spacing 2.5 Å) to broad long-range coverage (Coarse: 10–120 Å, average spacing 22 Å), with intermediate configurations bridging the two extremes. Within each configuration, the thresholds are spaced to provide approximately uniform coverage of the corresponding distance range. The effect of these configurations on prediction performance is reported in Section 2.3.

**Supplementary Table 1.** Distance threshold configurations.

| Configuration | Distance Thresholds (Å) | Range (Å) | Avg. Spacing (Å) |
|---|---|---|---|
| Baseline | — | — | — |
| Compact | 6.5, 7.5, 9.0, 11.5, 14.5, 19.0 | 6.5–19 | 2.5 |
| Near-range | 6.0, 8.0, 10.0, 12.0, 15.0, 20.0 | 6–20 | 2.8 |
| Dense | 7.0, 8.5, 10.0, 12.5, 16.0, 22.0 | 7–22 | 3.0 |
| Moderate | 7.0, 9.0, 11.0, 14.0, 18.0, 25.0 | 7–25 | 3.6 |
| Balanced | 8.0, 10.0, 15.0, 20.0, 30.0, 50.0 | 8–50 | 8.4 |

| | | | |
|---|---|---|---|
| Extended | 8.0, 12.0, 18.0, 25.0, 40.0, 65.0 | 8–65 | 11.4 |
| Broad | 10.0, 15.0, 22.0, 35.0, 55.0, 85.0 | 10–85 | 15.0 |
| Sparse | 8.0, 15.0, 25.0, 40.0, 60.0, 90.0 | 8–90 | 16.4 |
| Coarse | 10.0, 20.0, 35.0, 55.0, 80.0, 120.0 | 10–120 | 22.0 |

*Each configuration defines six distance thresholds (in Å) that partition the protein–RNA distance space into seven bins. Configurations are ordered by maximum threshold, from compact near-range sampling to coarse broad-range sampling.*

**Supplementary Note 2: Hyperparameter sensitivity analysis**

To assess the sensitivity of iSCALE to key architectural hyperparameters, we conducted a systematic search over five configurations varying hidden dimension, number of SSD layers, state dimension, convolution kernel size, and chunk size. All experiments used the protein–RNA binding $\Delta\Delta G$ dataset (S788) under five-fold cross-validation with random splitting.

**Supplementary Table 2.** Hyperparameter sensitivity of iSCALE on the protein–RNA binding $\Delta\Delta G$ task.

| Hidden Dim | Layers | State Dim | Conv Dim | Chunk Size | PCC | RMSE |
|---|---|---|---|---|---|---|
| 64 | 2 | 32 | 4 | 32 | **0.9661** | 0.467 |
| 64 | 3 | 32 | 4 | 32 | 0.9653 | **0.466** |
| 64 | 2 | 32 | 4 | 64 | 0.9649 | 0.469 |
| 128 | 3 | 16 | 4 | 64 | 0.9645 | 0.470 |
| 64 | 2 | 16 | 2 | 32 | 0.9642 | 0.471 |

*PCC: Pearson correlation coefficient; RMSE: root mean square error. Best values in each column are in bold.*

Across all configurations, PCC ranges from 0.964 to 0.966 and RMSE from 0.466 to 0.471, indicating that the prediction performance is stable with respect to these architectural choices. The final model adopts the configuration in the second row (hidden dimension 64, 3 SSD layers, state dimension 32, convolution kernel size 4, chunk size 32), which achieves the lowest RMSE while maintaining a competitive PCC.

To evaluate the contribution of the auxiliary spatial distribution prediction task, we varied the auxiliary loss weight $\boldsymbol{\lambda}$ across five values (Supplementary Table 3).

**Supplementary Table 3.** Effect of auxiliary task weight on prediction performance.

| Aux Weight | PCC |
|---|---|
| 0.2 | **0.9689 ± 0.0163** |
| 0.1 | 0.9608 ± 0.0311 |
| 0.0 | 0.9590 ± 0.0389 |
| 0.8 | 0.9655 ± 0.0239 |
| 1.0 | 0.9571 ± 0.0388 |

*Mean ± standard deviation across five folds. λ=0 corresponds to removing the auxiliary task entirely (equivalent to iSCALE*)*

The best performance is achieved at λ=0.2, with both the highest mean PCC and the lowest variance. Setting λ=0 (no auxiliary task) reduces PCC by 0.010 and increases variance, confirming that the auxiliary spatial distribution prediction provides a measurable regularization effect. Performance degrades slightly at λ=1.0, where the auxiliary loss dominates the total objective. The value λ=0.2 was used in all subsequent experiments reported in the main text.

#### Supplementary Note 3: Algorithm Implementation

Algorithm 1 summarizes the complete computational flow of iSCALE. $\mathrm{SSD}_1$ denotes the l-th bidirectional SSD layer (Section 4.5), and the same layers are reused across the three passes.

**Algorithm 1** iSCALE

**Require:** Wild-type sequence $P_{wt}$, mutant sequence $P_{mt}$, complex coordinates $\mathcal{S}$ (protein residue and binding-partner coordinates)

**Require:** Distance thresholds $\boldsymbol{\tau} = \{8,10,15,20,30,50\}$Å, block size $B = 32$, number of layers $L = 3$, auxiliary weight $\lambda = 0.2$

**Ensure:** Binding affinity change ΔΔGbind

▷ **Multi-scale Feature Construction**

$\mathbf{c}_{wt}, \mathbf{c}_{mt} \leftarrow$ ComputeMultiScaleCoupling $(P_{wt}, P_{mt}, \mathcal{S}, \boldsymbol{\tau})$

$\mathbf{S}_{wt} \leftarrow [\mathbf{x}_{seq,wt} \parallel \mathbf{x}_{evol,wt} \parallel \mathbf{c}_{wt}]$

$\mathbf{S}_{mt} \leftarrow [\mathbf{x}_{seq,mt} \parallel \mathbf{x}_{evol,mt} \parallel \mathbf{c}_{mt}]$

▷ **Three shared-parameter passes through the bidirectional SSD stack**

$\mathbf{H}_0^{(f)} \leftarrow \mathbf{S}_{wt} \oplus \mathbf{S}_{mt};\ \ \mathbf{H}_0^{(b)} \leftarrow \mathbf{S}_{mt} \oplus \mathbf{S}_{wt};\ \ \mathbf{H}_0^{(\mathrm{w})} \leftarrow \mathbf{S}_{wt}$

for $\ell = 1$ to L do

  $\mathbf{H}_\ell^{(f)} \leftarrow \mathrm{SSD}_\ell(\mathbf{H}_{\ell-1}^{(f)})$ ▷ forward concatenation (main task)

  $\mathbf{H}_\ell^{(b)} \leftarrow \mathrm{SSD}_\ell(\mathbf{H}_{\ell-1}^{(b)})$ ▷ reverse concatenation (main task)

$\mathbf{H}_{\ell}^{(\mathrm{w})}, \mathbf{S}_{blocks,\ell}^{(b)} \leftarrow \mathrm{SSD}_{\ell}(\mathbf{H}_{\ell-1}^{(\mathrm{w})})$ ▷ wild-type pass; block states (auxiliary)

for $b = 1$ to $N_{blocks}$ do

$\mathbf{s}_b \leftarrow \frac{1}{|\mathcal{D}_b|}\sum_{d\in\mathcal{D}_b} \mathbf{S}_{blocks}\,[b,d]$ ▷ Block representation

$\hat{\mathbf{c}}_b^{\mathrm{dist}} \leftarrow \mathrm{Softmax}(\mathrm{MLP}_{\mathrm{dist}}(\mathbf{s}_b))$ ▷Coupling distribution

$\hat{c}_b^{\mathrm{int}} \leftarrow \sigma(\mathrm{MLP}_{\mathrm{int}}(\mathbf{s}_b))$ ▷Coupling intensity

**end for**

$\mathcal{L}_{\mathrm{aux}} \leftarrow \frac{1}{N_{blocks}}\sum_{b=1}^{B}\left[\mathrm{KL}\left(\mathbf{c}_b^{\mathrm{dist}}, \hat{\mathbf{c}}_b^{\mathrm{dist}}\right) + \mathrm{BCE}\left(c_b^{\mathrm{int}}, \hat{c}_b^{\mathrm{int}}\right)\right]$

**end for**

$\mathcal{L}_{\mathrm{aux}} \leftarrow \mathcal{L}_{\mathrm{aux}}/\mathrm{L}$

▷ **Main task: differential conformational-coupling feature**

$\Delta_{coupling} \leftarrow \mathrm{GlobalPool}(\mathbf{H}_{mt}^{(f)}) - \mathrm{GlobalPool}(\mathbf{H}_{wt}^{(b)})$

$\widehat{\Delta\Delta G}_{bind} \leftarrow \mathrm{MLP}_{ddg}(\Delta_{coupling})$

▷ **Total loss**

$\mathcal{L}_{\mathrm{main}} \leftarrow \mathrm{MSE}(\widehat{\Delta\Delta G}_{bind}, \Delta\Delta G_{bind})$

$\mathcal{L}_{\mathrm{total}} \leftarrow \mathcal{L}_{\mathrm{main}} + \lambda \cdot \mathcal{L}_{\mathrm{aux}}$

**return** $\widehat{\Delta\Delta G}_{bind}, \mathcal{L}_{total}$
